# Enhancing Colorimetric LAMP Amplification Speed and Sensitivity with Guanidine Chloride

**DOI:** 10.1101/2020.06.03.132894

**Authors:** Yinhua Zhang, Guoping Ren, Jackson Buss, Andrew J. Barry, Gregory C. Patton, Nathan A. Tanner

## Abstract

Loop-mediated isothermal amplification (LAMP) is a versatile technique for detection of target DNA and RNA, enabling rapid molecular diagnostic assays with minimal equipment. The global SARS-CoV-2 pandemic has presented an urgent need for new and better diagnostic methods, with colorimetric LAMP utilized in numerous studies for SARS-CoV-2 detection. However, the sensitivity of colorimetric LAMP in early reports has been below that of the standard RT-qPCR tests, and we sought to improve performance. Here we report the use of guanidine hydrochloride and combined primer sets to increase speed and sensitivity in colorimetric LAMP, bringing this simple method up to the standards of sophisticated technique and enabling accurate and high-throughput diagnostics.

## Introduction

Loop-mediated isothermal amplification (LAMP) was developed as a simple nucleic acid amplification technique that reliably detects target sequence at a single reaction temperature without the need for sophisticated thermal cycling equipment (1). Advances in various detection technologies have helped establish LAMP as a useful and versatile tool for molecular diagnostics (2). Previously we reported a reliable visual detection method based on pH change during LAMP (3), enabling simple and low-cost applications suitable for point of care or field diagnostics. Colorimetric LAMP has been used in a range of applications: a large scale survey of Wolbachia-containing mosquitos (4); Grapevine red blotch virus without DNA extraction (5); testing urine samples for Zika virus (6) for example. In addition to use at the point of need, this detection method is amenable to medium-to high-throughput screening of large amounts of samples.

The recent and ongoing pandemic caused by SARS-CoV-2 has created an urgent demand for molecular diagnostics, requiring rapid creation of accurate and sensitive tests at unprecedented scale. LAMP presents an attractive option for diagnostic testing due to compatibility with simple colorimetric detection and relatively unpurified sample input and several studies have demonstrated its usage in diagnostics of SARS-CoV-2 (7–13) including potential utility for both simple, rapid testing and higher throughput applications (11, 13). To date these studies have demonstrated excellent specificity with LAMP, however, sensitivity has generally been lower than typical RT-qPCR assays (e.g. 87.5% sensitivity as compared to a standard RT-qPCR test) (14).

Accordingly, we set out to improve the sensitivity of RT-LAMP, screening new and published primer sets for SARS-CoV-2 RNA detection and identified two sets with marked increase in sensitivity. We also screened many compounds and reaction conditions for improving LAMP speed and detection sensitivity, with the addition of guanidine hydrochloride providing a notable enhancement to both. Here we describe these studies and the performance improvement resulting from use of guanidine and the combination of primer sets together in one LAMP reaction. Additionally, use of absorbance measurement of the color change of the pH sensitive dye phenol red enabled more sensitive detection of low-copy amplification, reliably calling positive amplification down to 10 copies of input RNA in 30 minutes. These modifications to standard RT-LAMP conditions will further the ability of LAMP to support sensitive molecular diagnostic tests, for the current COVID crisis and future diagnostic needs.

## Results

In order to improve the amplification in RT-LAMP reactions, we screened a range of compounds and additives commonly used to enhance PCR or other isothermal methods (e.g. DMSO, poly(ethylene glycol) compounds, betaine, single-stranded DNA binding proteins). While most of these compounds had no, very little, or negative effects on LAMP, we found that guanidine chloride showed dramatic improvement of speed on LAMP (Fig. 1A). Other guanidine containing compounds such as guanidine thiocyanate and arginine were also found to have a stimulatory effect, but with a narrower workable range and accordingly we focused on guanidine hydrochloride. We determined that an optimal range of guanidine chloride concentration for 3 primer sets used in SARS-CoV-2 RNA detection and all showed the greatest stimulation at around 40–50 mM when used with the colorimetric LAMP master mix (Fig. 1B), indicating the stimulation is likely general to colorimetric LAMP and not specific to a particular primer set.

**Figure 1.**
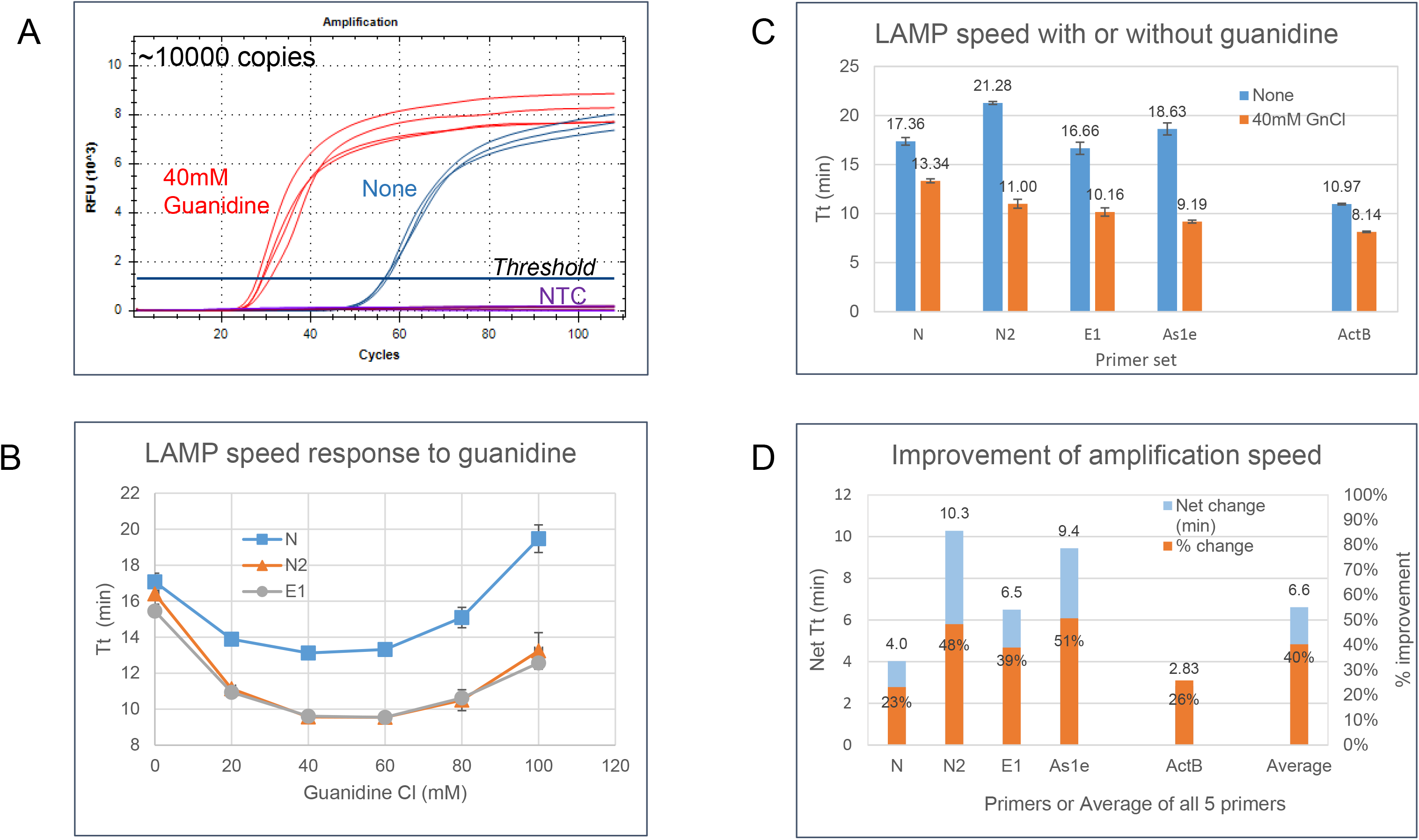
Increase in LAMP amplification speed by addition of guanidine chloride. A. Real-time fluorescence curves showing increase of amplification speed by including guanidine chloride in the LAMP reaction. Speed was determined by the time the signal crossing a threshold line (Tt, time to threshold) automatically set by the software. B. Determine optimal guanidine concentration. The speed of three primer sets (N, N2 and E1) were plotted with averages of 4 repeats. C. Amplification speed of 5 primer sets with or without guanidine chloride. There were ~10000 copies synthetic covid-19 RNA primers for N, N2, E1 and As1e primers. For ActB primer set, 1.0 Jurkat total RNA was used in the amplification. D. Comparing stimulation of LAMP speed by guanidine chloride. The primary Y axis shows the net change of Tt between without and with guanidine. The secondary axis shows the percentage change (net Tt/Tt of no guanidine)

Using this optimal concentration of guanidine, we checked the speed enhancement with four SARS-CoV-2 RNA primer sets (two for Gene N (N, N2), one for Gene E (E1) and a published set for Orf1a (As1e)) using 10,000 copies of synthetic SARS-CoV-2 RNA template and one primer set for human beta actin (ACTB), often used as an extraction control assay for patient samples, with 1 ng of total Jurkat RNA template (Fig 1C-D). Addition of guanidine increased LAMP speed by nearly 2-fold for three of the primer sets: 21.3 minutes (threshold time from CFX-96 instrument) to 11 minutes for the N2 primer set; 16.7 min to 10.2 for the E1 primer set; and 18.3 minutes to 9.2 minutes for the As1e primer set. The N primer set showed less stimulation (17.4 to 13.3 minutes) as did the already very fast ACTB primer set (11 to 8.14 minutes). On average together for all five primer sets, guanidine chloride shortened the time to threshold detection by 6.6 minutes or ~40%. Importantly, this increase in detection speed did not cause any increase in no-template control amplification with any of the five primer sets (Fig. 1A, Fig. 2C-D), indicating the stimulation by guanidine is specific to the interaction between primers and their intended templates.

**Figure 2.**
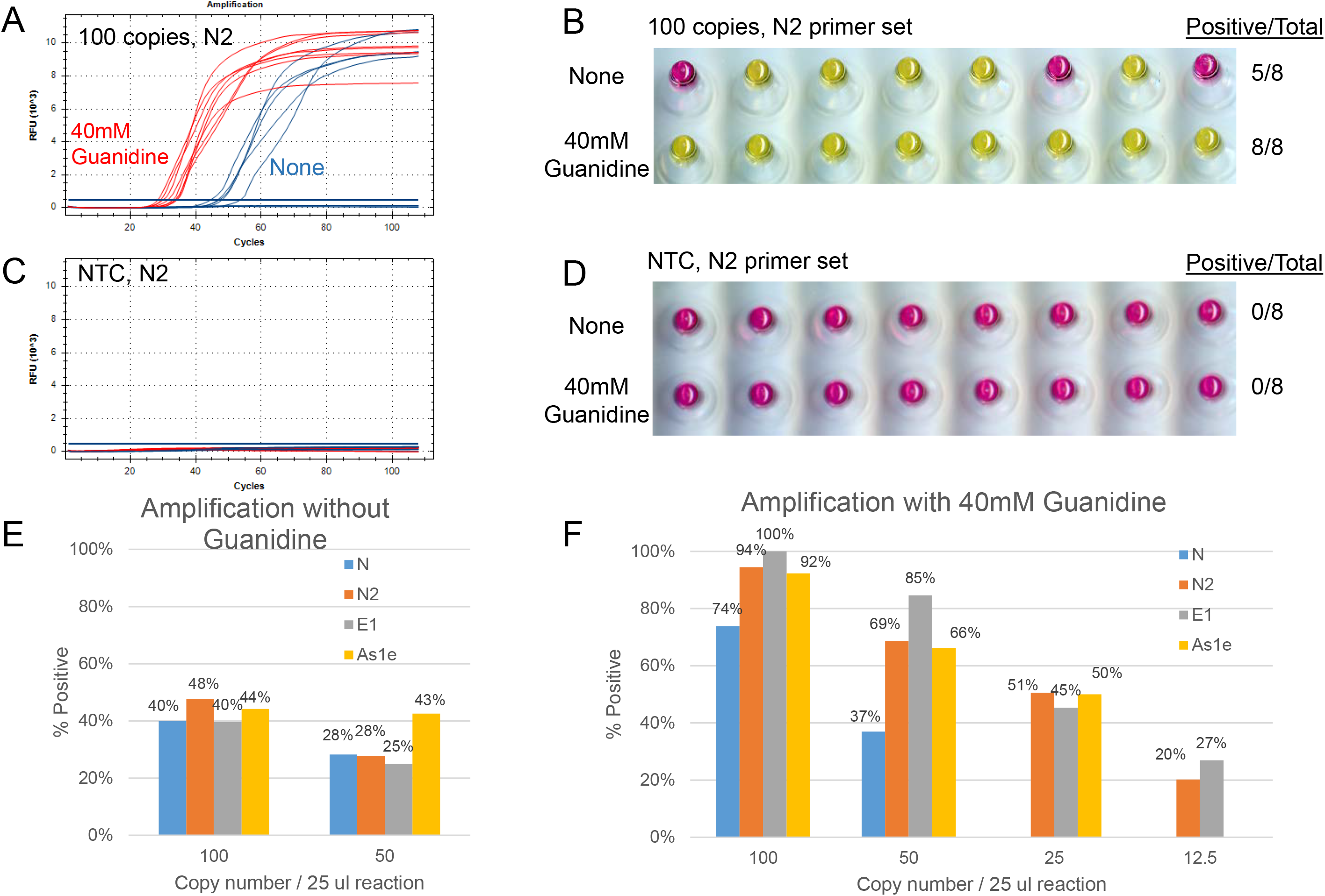
Enhancing detection sensitivity with guanidine chloride. A. Real time curve of amplifying 100 copies of SARS-CoV-2 RNA using N2 primers with or without guanidine, replicates of 8 reactions each. B. End point color change scoring of positive amplification of panel A. C. No template control in real time monitoring. D. No template control with end point color scoring of panel C. E. Percentage amplification without guanidine with 100 and 50 copies of template RNA. F. Percentage amplification in the presence of 40mM guanidine with 100, 50, 25 and 12.5 copies of template RNA.

Having observed such a significant effect on reaction speed, we next examined whether guanidine also has an effect on LAMP amplification sensitivity using lower RNA template inputs. By monitoring the reaction in real time, we found that guanidine also significantly shortened the reaction time with low template input (Fig. 2A) at a similar degree to that with high template input observed above. For scoring positive amplification, we relied on end point scoring based on the color change from pink to yellow (Fig. 2B), with results concordant with real time observation. Multiple reactions were performed for four primer sets using 100 or 50 copies of synthetic SARS-CoV-2 RNA. In all cases, guanidine significantly improved the percentage of positive detection. With 100 copies of RNA template, the detection percentage increased from just below 50% without guanidine to over 90% when it was added to reactions with the N2, E1 and As1e primer sets, with primer set N showing increase from 40% to 74% (Fig. 2E). For 50 copies, the percentage increased from lower 30% to around 70% (Fig. 2F) for sets N2, As1e, and E1. Overall, in the presence of guanidine, it is possible to successfully amplify for about 50% for 25 copies of input, slightly higher than that for 100 copies without guanidine. This accordingly indicates an increase in sensitivity of approximately 4-fold by including 40 mM guanidine hydrochloride in the LAMP reaction.

As each primer set was designed to detect different regions of the SARS-CoV-2 RNA sequence, combining two primer sets could potentially increase detection if there is no interference of the two primer sets. We tested this idea in pair wise combinations of the most sensitive LAMP primer sets (N2, E1, As1e) as well as using the three primer sets together. In these combination reactions, the concentration of each primer set was kept the same as in the reaction with only a single set, so the total primer concentration is doubled and tripled respectively. Based on the real time curves of these reactions, in the presence of guanidine, amplification in double and triple primer reactions started much earlier than without guanidine (Fig.3A). Almost all positive reactions started at a similar early timepoint, while those without guanidine initiated at scattered times much later. For the end point color change scoring, the color changed from pink to yellow completely in the presence of guanidine while the color change is only partial in no guanidine reactions (Fig. 3B), reflecting the slower amplification.

**Figure 3.**
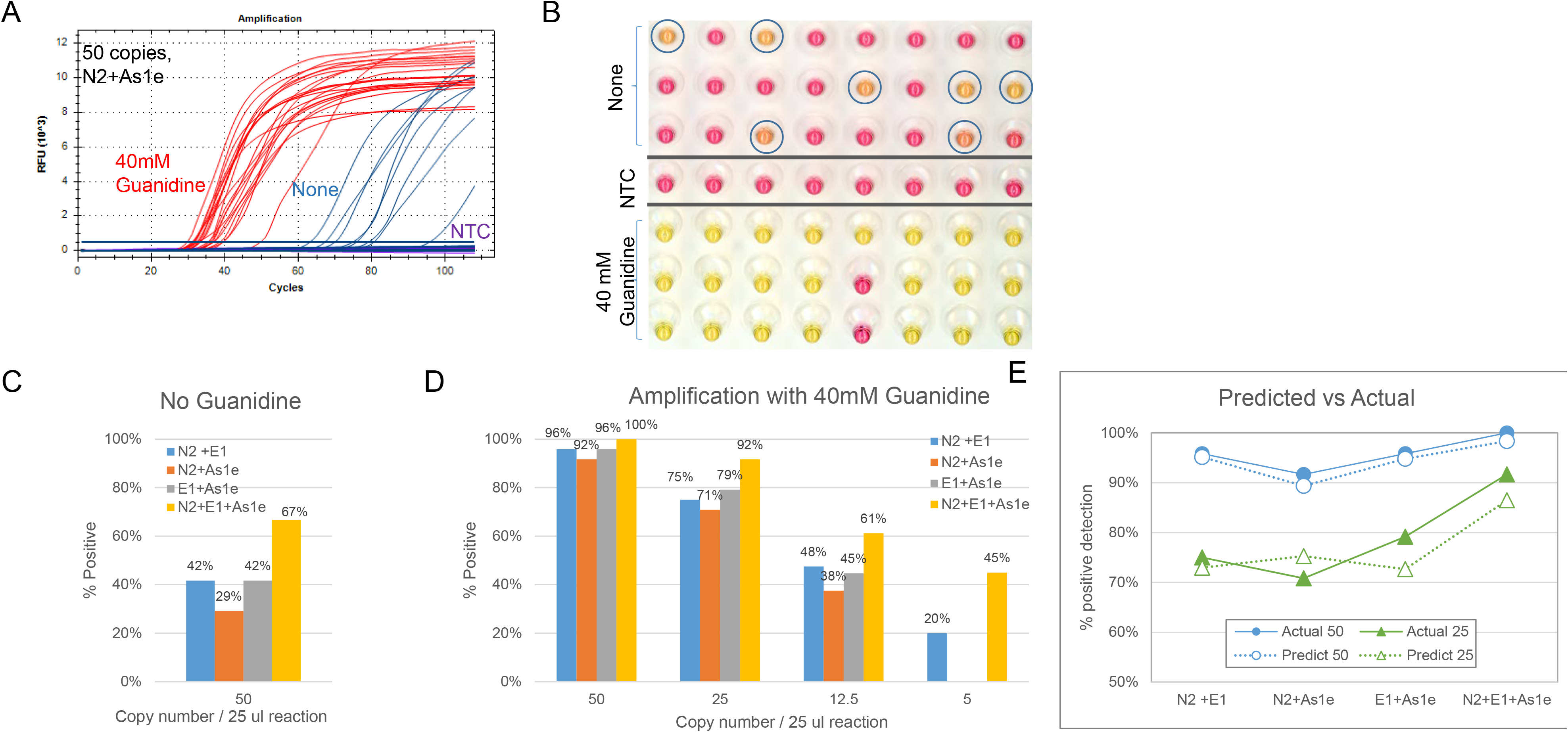
Guanidine enables high sensitivity detection by combining two or three primer sets. A. Real time curves of 24 reactions each with (red curves) or without (blue curves) guanidine using 50 copies of COVID RNA template and N2+As1e primer sets. B. End point color change scoring for positive amplification in panel A. In the presence of guanidine, positive reactions changed to yellow completely while in no guanidine reactions the color change is only partially changed (circled). C. Positive amplification rate with dual and triple primer combinations with N2, E1 and As1e primers without guanidine with 50 copies of template. D. Positive amplification rate with dual and triple primer combinations with N2, E1 and As1e primer sets in the presence of 40mM guanidine chloride with 50, 25, 12.5 and 5 copies of template. E. Actual detection rate correlates strongly with predicted frequency

In the absence of guanidine, the detection rate with double primers and 50 copies of RNA template only showed a slight increase, from just below 30% to 40% (Fig. 2C and 3C). The positive rate for triple primer set was higher, measured at 67%. When guanidine was added, all combinations showed a positive rate of over 92% with the triple primer set at 100% (Fig 3D). When even lower templates were tested, 25 copies were detected nearly 75% of the time with dual primer combinations and 92% with the triple primer reactions. With 12.5 copies this rate was 45% for dual and 61% triple-primer reactions, 5 copies 20% and 45%. In all of the dual and triple primer combination tests, there was no signal in NTC reactions when incubated to 40 minutes (Fig. 3A-B). Given the improvement in positive detection rate, we estimate dual primer combination in the presence of guanidine can almost double the detection frequency of SARS-CoV-2 RNA, and more than doubled with all three primers.

The increased detection frequency using dual or triple primers seemed to follow a simple combined detection probability. Based on the detection frequency of 50 or 25 copies of single primer set (Fig. 2F), theoretical combined detection probabilities of all combinations were calculated assuming each primer set detects its target independently and there is no interference between any combination. These predicted numbers strongly correlate with the actual detected frequency (Fig. 3D) for both 50 and 25 copies (Fig. 3E). We also tested whether the increased detection is due to a simple primer concentration increase. When the concentration of a single primer set (N2) was increased to 2- or 3-fold to match the total primer concentration of the dual- and triple-primer reactions, we found there was no increased detection frequency (data not shown). However, there was noticeable increase of NTC signal. Thus, increased sensitivity is not due to simply more primer being available, but rather more due to LAMP reactions of multiple primer sets occurring independently at the same time, and effectively increasing the available number of templates for the same target.

Next we sought to improve detection sensitivity and evaluate high-throughput compatibility by quantitatively detecting the colorimetric change of our reactions using spectrophotometric measurements. We combined the optimal guanidine hydrochloride concentration (40mM) with the dual primer sets (N2 and E1) described above and performed colorimetric LAMP reactions with a 2-fold dilution series of synthetic COVID-19 RNA. Using the ratio of absorbance measurements of the two phenol red peaks (yellow Abs_432_/ pink Abs_560_) as the primary metric, we observed a positive correlation with copy number (Figure 4). Applying a threshold of 0.704, equivalent to the 99.7% confidence interval (μ_NTC_ + 3σ_NTC_) of NTC (n=8), we determined that 100% of all samples containing 20-160 copies rxn^−1^ tested positive. Notably, 62.5% of samples containing only 10 copies rxn^−1^ also tested positive, illustrating a marked improvement on sensitivity. These reactions were incubated for only 20 minutes, with the more sensitive instrumented measurement enabling calling positives much earlier than detection simply by eye.

**Figure 4.**
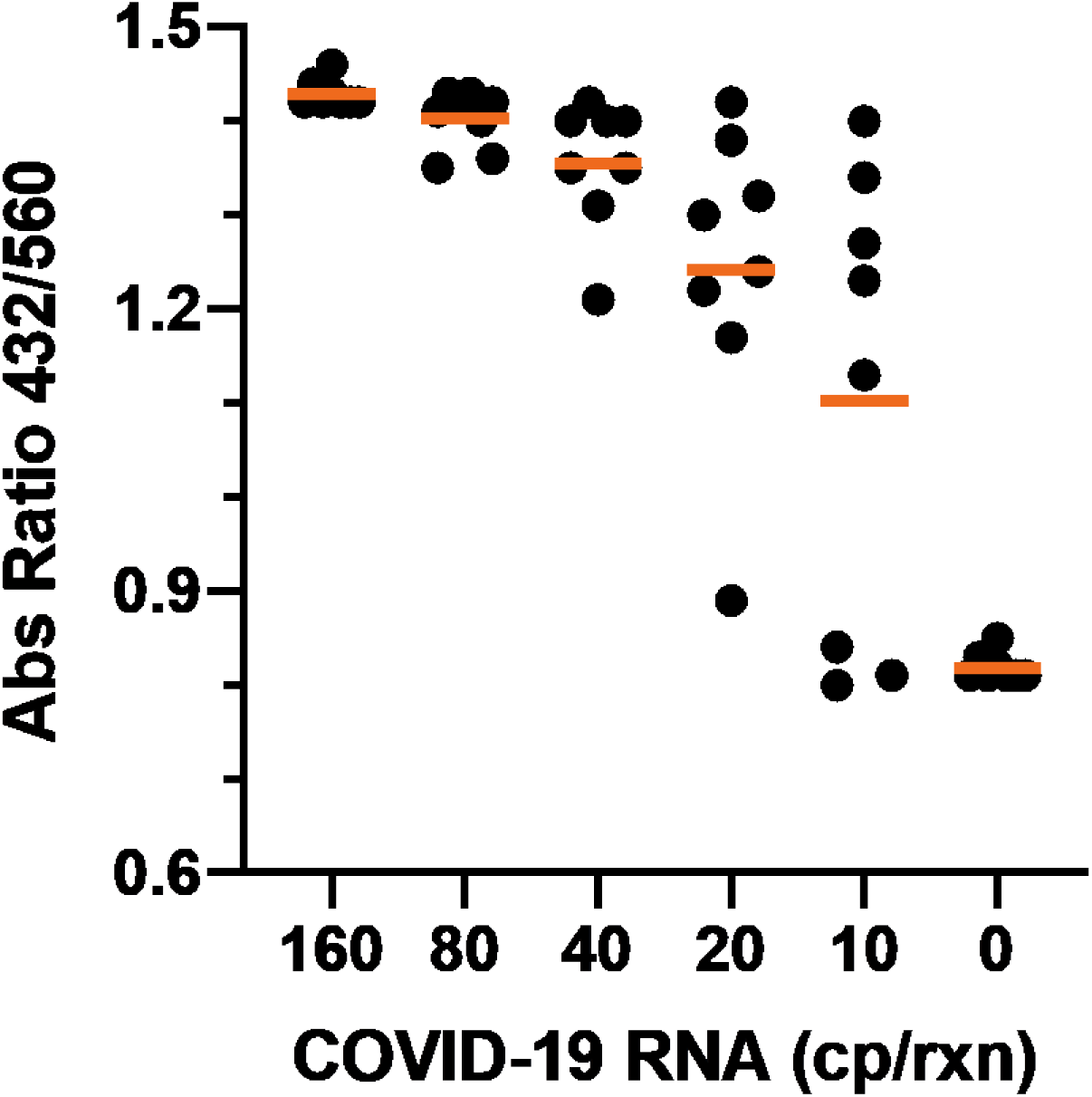
Endpoint Absorbance Measurements of Optimized Colorimetric LAMP on Synthetic COVID-19 RNA. Colorimetric LAMP was performed on a synthetic COVID-19 RNA (TWIST; 160, 80, 40, 20, 10, and 0 copies rxn^−1^) at 65°C for 20 min and endpoint absorbance at 432nm and 560nm was determined after a rapid cooling step. The ratio of the two absorbance readings (Abs_432_/Abs_560_) is plotted against the corresponding RNA copy number (n = 8). The mean for each group is displayed in orange. Results indicate a positive correlation between absorbance ratio and copy number (μ ± (J: 160cp – 1.410 ± 0.017; 80cp – 1.383 ± 0.084; 40cp – 1.321 ± 0.084; 20cp – 1.190 ± 0.200; 10cp – 1.015 ± 0.298; NTC – 0.679 ± 0.008), and suggest that simple, endpoint spectrophotometric analyses provide a rapid and sensitive approach for the detection of COVID-19 RNA by Colorimetric LAMP.

## Discussion

We present here a significant increase in the speed and sensitivity of RT-LAMP reactions by addition of guanidine hydrochloride. This effect is compounded by additional enhancement of combining two or three LAMP primer sets, enabled by guanidine and resulting in additional improvement in sensitivity without increase in nonspecific amplification. These enhanced reactions also make it possible to utilize shorter detection times and better discrimination of color change with the colorimetric LAMP detection based on pH-sensitive dye.

This report describes the enhancement of RT-LAMP by addition of guanidine chloride, yet the mechanism of its action is yet to be elucidated. Based on its consistent enhancement of reactions with different primer sets, we speculate that guanidine enhances the base pairing between primers and their target sequences. Consistent with this interpretation, it also significantly shortens the reaction time for helicase-dependent amplification (data not shown). It is unlikely due to the modulation of enzyme activity, as additional enzyme, either reverse transcriptase or *Bst* 2.0 DNA polymerase, did not have such an effect. While the primary focus here is RT-LAMP reactions, the guanidine enhancement is equally effective with LAMP using DNA inputs (data not shown).

Measurement with an absorbance plate reader also demonstrated a notable increase in reaction sensitivity, clearly calling reaction positives that would have been indeterminate at best by visual detection. While visual readout is well-suited to simple field and point-of-care applications, endpoint reading of plates by absorbance provides a quantitative data record more amenable to high-throughput settings. Plates (96- or 384-well) could be incubated in simple heating chambers and endpoint absorbance used for analysis, and if paired with liquid handling plate filling a large number of samples could be processed with the simple 20–40 minute heating at 65 °C and endpoint plate read. One example of this workflow is shown in Figure 5, with automated liquid handling and nucleic acid extraction for plate setup, followed by heating in an oven and measurement of absorbance to determine presence/absence of target. An Emergency Use Authorization was recently granted to Color Genomics for a SARS-CoV-2 test much as laid out here, proving the principle as a potential high-throughput workflow. Taken together, the improvements to LAMP presented here describe a significant increase in sensitivity for this powerful isothermal method to be more fully utilized for molecular diagnostics.

**Figure 5.**
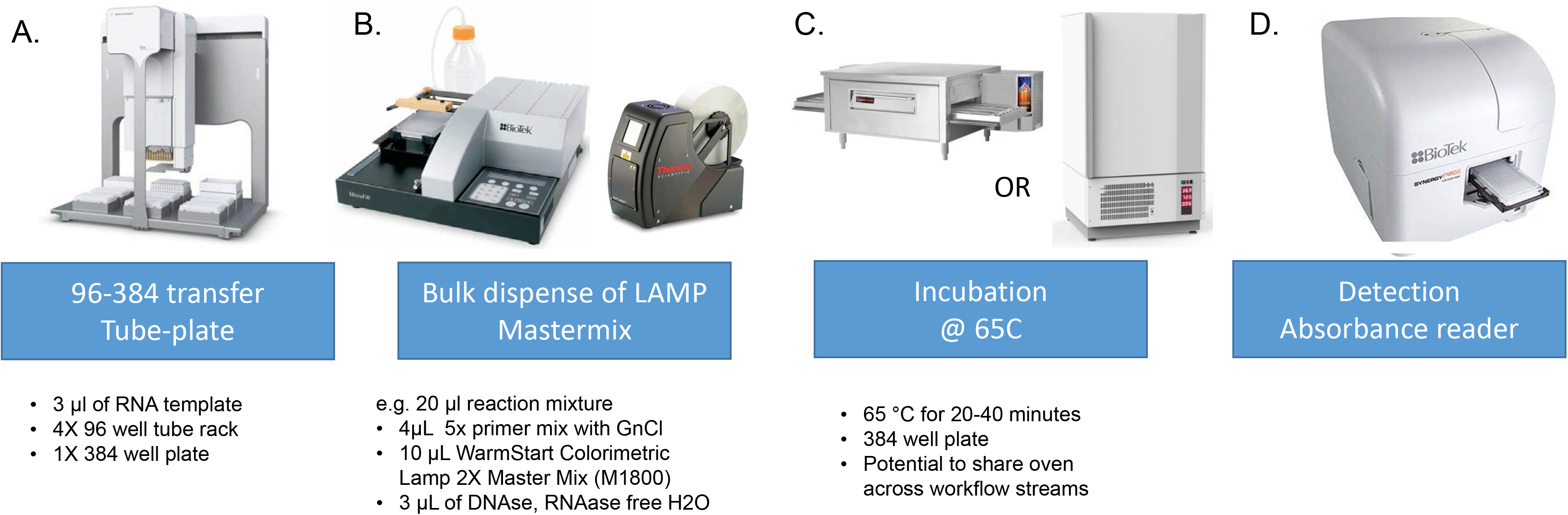
Example of an automated workflow supporting high-throughput endpoint detection of colormetric LAMP. A. 96 channel liquid handler (Agilent Bravo, Santa Clara, CA, P/N G5563A) for transfer of extracted nucleic acids into 384-well absorbance plate. B. Bulk dispense of LAMP Mastermix (BioTek Ufill, AF1000A) into 384-well plates and plate sealing (ALPS AB-3000 Plate Sealer ThermoFisher, Waltham, MA). C. Isothermal incubation of sealed plates using a custom conveyor oven (Intek Custom Electric Conveyer Oven, Union, Missouri), or automated random-access incubator (Liconic STR240, Mauren, Liechtenstein). D. Endpoint detection using absorbance plate reader (BioTek Synergy Neo2, Winooski, VT).

## Materials and Methods

LAMP primers targeting different regions of SARS-CoV-2 sequence (GenBank accession number MN908947) were designed using the online software Primer Explorer V5 (https://primerexplorer.jp/e/). Primers N (14) and As1e (12) appeared in previous preprints. We screened 7 additional new primer sets and found that N2 and E1 gave best performance. Primer sequences are listed in Table 1. Oligos were synthesized at Integrated DNA Technologies with standard desalting. Synthetic COVID-19 RNA containing equal ratio of most viral genome regions was purchased from Twist Bioscience (Twist Synthetic SARS-CoV-2 RNA Control 2 (MN908947.3) - SKU: 102024), which came at a concentration of 1 million copies per ul. The RNA was diluted to lower concentrations in 10ng/ul Jurkat total RNA and their concentration were estimated using RT-qPCR with primer and probe sets for CDC N1 and Charité/Berlin Gene E. RT-LAMP reactions were performed using WarmStart® Colorimetric LAMP 2X Master Mix (DNA & RNA) (M1800) supplemented with 1 μM SYTO®-9 double-stranded DNA binding dye (Thermo Fisher S34854) and incubated on a real-time qPCR machine (BioRad CFX96) for 107 cycles with signal acquisition for every 15 seconds (total incubation time ~40 min). The color of the finished reactions was recorded using an office flatbed scanner. Guanidine chloride solution was made with Sigma RDD001 and adjusted to pH ~8 with KOH before adding to the colorimetric LAMP reaction.

**Table 1.**
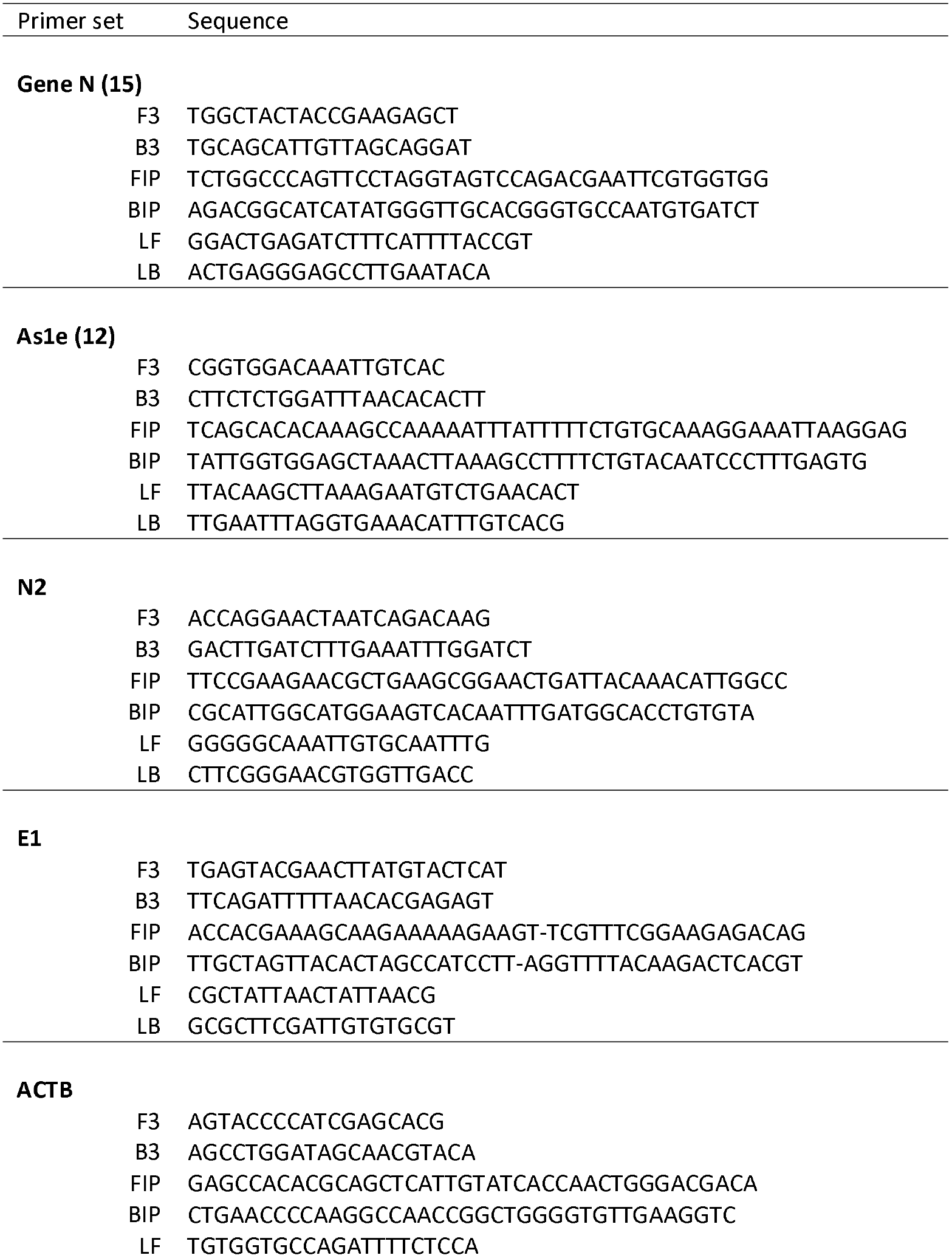
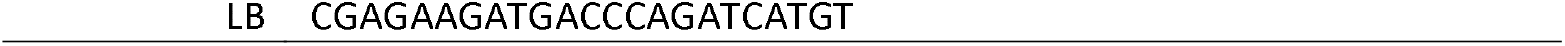
LAMP primer sequences and source

For plate reader absorbance measurement we applied an ABI MicroAmp seal to the Bio-Rad HSP9601 microplate and incubated the reactions at 65°C for 20 min in a ThermoMixer C equipped with a ThermoTop heated lid and a 96-well adaptor. We then rapidly cooled the sample to reduce condensation effects and read absorbance [SpectraMax M5; BioTek Synergy Neo2] at 432nm (yellow) and 560nm (red), corresponding to the pH-dependent maxima of Phenol Red.

